# Polygenic Hyperlipidemias and Coronary Artery Disease Risk

**DOI:** 10.1101/735167

**Authors:** Pietari Ripatti, Joel T Rämö, Nina J Mars, Sanni Söderlund, Christian Benner, Ida Surakka, Tuomo Kiiskinen, Aki S Havulinna, Priit Palta, Nelson B Freimer, Veikko Salomaa, Matti Pirinen, FinnGen Aarno Palotie, Marja-Riitta Taskinen, Samuli Ripatti

**Author notes:** Corresponding author Street address: Tukholmankatu 8, FI-00290 Helsinki, Finland, Correspondence: P.O. Box 20, FI-00014 University of Helsinki, Helsinki, Finland. These authors contributed equally to this work (Marja-Riitta Taskinen, Samuli Ripatti). Journal Subject Terms: Biomarkers, Lipids and Cholesterol, Epidemiology, Risk Factors, Genetics, Coronary Artery Disease.

## Abstract

**Background:** Hyperlipidemia is a highly heritable risk factor for coronary artery disease (CAD). Monogenic familial hypercholesterolemia associates with higher increase in CAD risk than expected from a single LDL-C measurement, likely due to lifelong cumulative exposure to high LDL-C. It remains unclear to what extent a high polygenic load of LDL-C or TG-increasing variants associates with increased CAD risk.

**Methods and Results:** We derived polygenic risk scores (PRS) with ∼6M variants for LDL-C and TG with weights from a UK biobank-based genome-wide association study with ∼500K samples. We evaluated the impact of polygenic hypercholesterolemia and hypertriglyceridemia to lipid levels in 27 039 individuals from the FINRISK cohort, and to CAD risk in 135 300 individuals (13 695 CAD cases) from the FinnGen project.

In FINRISK, LDL-C ranged from 2.83 (95% CI 2.79-2.89) to 3.80 (3.72-3.88) and TG from 0.99 (0.95-1.01) to 1.52 (1.48-1.58) mmol/l between the lowest and highest 5% of the respective PRS distributions. The corresponding CAD prevalences ranged from 8.2% to 12.7% for the LDL-C PRS and from 8.2% to 12.1% for the TG PRS in FinnGen. Furthermore, CAD risk was 1.36-fold higher (OR, 95% CI 1.24-1.49) for the LDL-C PRS and 1.31-fold higher (1.20-1.44) for the TG PRS for those with the PRS >95^th^ percentile vs those without. These estimates were only slightly attenuated when adjusting for a CAD PRS (OR 1.26 [95% CI 1.15-1.39] for LDL-C and 1.21 [1.10-1.32] for TG PRS).

**Conclusions:** The CAD risk associated with a high polygenic load for lipid-increasing variants was proportional to their impact on lipid levels and mostly independent of a CAD PRS. In contrast with a PRS for CAD, the lipid PRSs point to known and directly modifiable risk factors providing more direct guidance for clinical translation.

## Introduction

Hypercholesterolemia, particularly high LDL-cholesterol (LDL-C), is an established, heritable, and treatable risk factor for coronary artery disease (CAD).^1, 2^ Additionally, accumulating evidence suggests that increased triglycerides (TG; hypertriglyceridemia) are causally linked to CAD.^3-5^

Increased levels of both LDL-C and TGs result from a combination of genetic and non-genetic factors.^6, 7^ Genetic factors include rare highly penetrant variants and a long tail of common variants with smaller effect sizes. While high impact variants in the *LDLR, PCSK9*, and *APOB* genes cause familial hypercholesterolemia, it has also been suggested that similarly high LDL-C levels could result from a high polygenic burden of LDL-C-increasing variants.^8, 9^ While monogenic FH with an identified mutation associates with a higher CAD risk than expected on the basis of a single LDL-C measurement, the contribution of an accumulation of a large number of LDL-C-increasing alleles to CAD risk is unclear.^10^

Similarly to hypercholesterolemia, both polygenic burden and highly penetrant variants contribute to hypertriglyceridemia.^6^ However, highly penetrant variants underlying hypertriglyceridemia are much fewer and very rare (estimated population prevalence 1:1 000 000).^6^ On the other hand, many individuals with hypertriglyceridemia have a high polygenic burden of TG-increasing variants.^6^ Unlike LDL-C, it is unknown whether genetically increased TGs confer higher CAD risk than non-genetic hypertriglyceridemia. Genetics supporting a causal link between hypertriglyceridemia and CAD, and the evidence for beneficial therapeutic reducing of TGs to reduce CVD risk, however, highlight the potential also for association between polygenic load of TG elevating alleles and CAD risk.^3-5, 11, 12^

In this cohort study of 27 039 individuals from the Finnish FINRISK population cohort with lipid measurements, and 135 300 individuals including 13 695 CAD cases from the FinnGen project, we evaluated the impact of high polygenic LDL-C and TG to CAD risk. We developed separate genome-wide PRSs for both LDL-C and TG to define polygenic hypercholesterolemia and hypertriglyceridemia. First, we tested to what extent PRSs for LDL-C and TG associate with measured lipid levels. Second, we tested to what degree polygenic hypercholesterolemia and polygenic hypertriglyceridemia associate with increased risk for CAD.

## Methods

### Ethics Statement

All samples were collected in accordance with the Declaration of Helsinki. For the Finnish Institute of Health and Welfare (THL) driven FinnGen preparatory project (here called FinnGen), all patients and control subjects provided informed consent for biobank research, based on the Finnish Biobank Act. Alternatively, older cohorts were based on study-specific consents and later transferred to the THL Biobank after approval by Valvira, the National Supervisory Authority for Welfare and Health. Recruitment protocols followed the biobank protocols approved by Valvira. The FinnGen project was additionally approved by THL (approval numbers THL/2031/6.02.00/2017, and amendments THL/341/6.02.00/2018, THL/2222/6.02.00/2018, and THL/283/6.02.00/2019). Written informed consent was obtained from all participants except the 1992 FINRISK survey, for which verbal informed consent was obtained as required by legislation and ethics committees at the time. Earlier FINRISK surveys were approved by various ethics committees.^13^ The Coordinating Ethics Committee of the Helsinki and Uusimaa Hospital District approved the FinnGen project (number HUS/990/2017) and the 2007 and 2012 FINRISK surveys. The North West Multi-Centre Research Ethics Committee approved the UKBB study.

### Subjects and Measurements

The National FINRISK Study is a Finnish population survey conducted every 5 years since 1972 with independent, random, and representative samples across the country.^13^ We used 27 039 individuals from the 1992 to 2012 collections. Circulating biochemical markers were measured from venous blood samples drawn after a minimum of 4-h fast using standard methods.^13^ The effect of lipid-lowering therapy in those using medication was adjusted for by dividing LDL-C by 0.7 as utilized previously.^10^ LDL-C was calculated using the Friedewald formula.^14^ Non-HDL-C was calculated as total cholesterol (TC) - HDL-C and remnant cholesterol (remnant-C) as TC - HDL-C - LDL-C.

The FinnGen preparatory phase aggregates Finnish biobank samples and currently comprises 135 300 participants.^15^ The samples have been linked with national hospital discharge and causes-of-death registries. Clinical CAD event endpoints were constructed from major adverse coronary events (MACE) defined as either myocardial infarction (MI) (International Classification of Diseases [ICD]-10 codes I20.0 or I21-22, ICD-9 410 or 411.0, or ICD-8 410 or 411.0 for hospital discharge; or ICD-10 I21-25, I46, R96, or R98, ICD-9 410-414 or 798 [excluding 7980A], or ICD-8 410-414 or 798 for main cause-of-death) or coronary revascularization (coronary angioplasty [PCI] or coronary artery bypass grafting [CABG]).^16^

The UK Biobank comprises extensive phenotypic data on some 500 000 individuals of the general UK population between 40 and 69 years.^17^ All participants were interviewed, answered standardised questionnaires, and had physical measurements taken at baseline. The UKBB cohort was linked with national Hospital Episode Statistics, cancer, and death registry data.^17^ Circulating biochemical markers were measured from serum samples drawn after a mean fasting time of 3.8 hours. LDL was measured using enzymatic selective protection and TG using enzymatic methods (http://biobank.ctsu.ox.ac.uk/crystal/crystal/docs/serum_biochemistry.pdf).

### Genotyping and Polygenic Risk Score Calculation

Samples were genotyped and imputed using standard methods as described in S1 Text.

The LDL-C, TG, and CAD PRSs were calculated as the sum of the risk allele dosages weighted by their effect sizes using LDpred.^18^ The recent LDpred method is a Bayesian approach to calculate a posterior mean effect size for each variant based on a prior of effect size and linkage disequilibrium (LD; a measure of how much a variant correlates with other variants).^18^ Whole-genome sequences from 503 European samples from the 1000 Genomes project phase 3 served as the LD reference population for LDpred.^19^ We utilised the infinitesimal prior on the fraction of causal variants in a given phenotype.

The weights for the lipid PRSs were based on a genome-wide association study (GWAS) of 468 732 samples with LDL-C measurements and 469 240 with TG measurements from the UKBB. As part of quality control, related subjects and subjects taking lipid-lowering medicine were excluded. We performed the lipid GWAS using BOLT-LMM and adjusted for sex, age and the first 15 principal components.^20^ The weights for the CAD PRS were based on summary statistics obtained from a GWAS of ischemic heart disease (IHD) in the UKBB (PheCode 411) performed using SAIGE.^21^ The PRSs were calculated using PLINK 2.0 Alpha 1.^22^ The final PRSs included 5 707 489 variants for LDL-C and TG and 5 709 394 variants for CAD.

### Statistical Analysis

Variation explained by PRSs was estimated as adjusted *r*^*2*^ from linear regression, with residual lipid measurements after adjusting for age and sex as the response. TG measurements were additionally log-transformed. Bootstrapping with percentile CIs of served to estimate median lipid levels in PRS bins. Binomial logistic regression served to estimate odds ratios (OR) for CAD outcomes. The logistic regression models were adjusted for age, sex, first ten principal components, and genotyping batch. All tests were two-sided. Statistical analyses were performed using R (version 3.6.1).^23^

## Results

### Polygenic Hyperlipidemias and Lipid Levels

We first defined PRSs for LDL-C and TG using an approach of reweighting the effects of genome-wide variants using GWAS summary statistics and the LD structure of a reference population implemented in the software package LDpred.^18^ As the largest freely available population-wide dataset of lipid measures and genetic markers, we drew the summary statistics from a GWAS of ∼500 000 individuals from the UKBB with lipid measures and tested the association between PRSs and lipid levels in the Finnish FINRISK study, independent of the original GWAS dataset. The FINRISK study comprised 27 039 individuals randomly drawn from the Finnish population (Table 1). Median LDL-C was 3.39 mmol/l and TG 1.19 mmol/l in the whole cohort with slightly lower values in the more recent collections (S1 Figure).

**Table 1.**
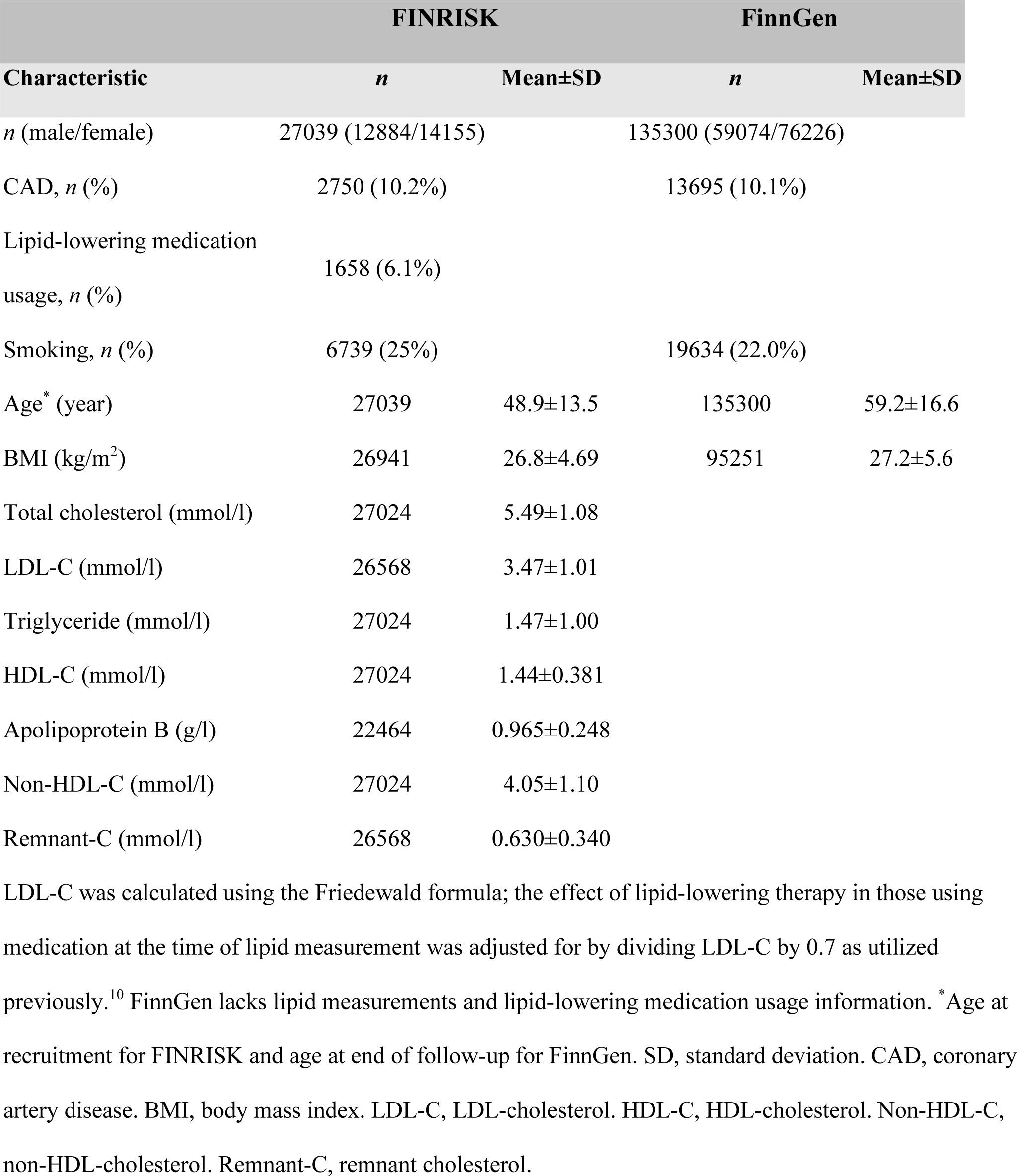
Clinical and Metabolic Characteristics of Individuals.

**Table 2.** CAD Prediction with Lipid and CAD PRSs.

| Predictors | AUC | OR (95% CI) | *p* |
| --- | --- | --- | --- |
| PRS<sub>LDL-C</sub> | 0.877 | 1.16 (1.13-1.18) | $< 2 \times 10^{-16}$ |
| PRS<sub>TG</sub> | 0.877 | 1.13 (1.10-1.15) | $< 2 \times 10^{-16}$ |
| PRS<sub>CAD</sub> | 0.881 | 1.33 (1.30-1.36) | $< 2 \times 10^{-16}$ |
| PRS<sub>CAD</sub> + PRS<sub>LDL-C</sub> | 0.881 |  |  |
| PRS<sub>CAD</sub> | | 1.32 (1.29-1.34) | $< 2 \times 10^{-16}$ |
| PRS<sub>LDL-C</sub> | | 1.12 (1.10-1.15) | $< 2 \times 10^{-16}$ |
| PRS<sub>CAD</sub> + PRS<sub>TG</sub> | 0.881 |  |  |
| PRS<sub>CAD</sub> | | 1.32 (1.29-1.35) | $< 2 \times 10^{-16}$ |
| PRS<sub>TG</sub> | | 1.08 (1.06-1.11) | $5.86 \times 10^{-13}$ |
| PRS<sub>CAD</sub> + PRS<sub>LDL-C</sub> + PRS<sub>TG</sub> | 0.881 |  |  |
| PRS<sub>CAD</sub> | | 1.31 (1.28-1.34) | $< 2 \times 10^{-16}$ |
| PRS<sub>LDL-C</sub> | | 1.10 (1.08-1.13) | $< 2 \times 10^{-16}$ |
| PRS<sub>TG</sub> | | 1.05 (1.03-1.08) | $1.28 \times 10^{-5}$ |

ORs and AUCs for CAD with continuous LDL-C, TG, and CAD PRSs as predictors estimated using logistic regression. All models were additionally adjusted for age and sex. AUC, area under the ROC curve. ROC, receiving operating characteristic. OR, odds ratio. CI, confidence interval. CAD, coronary artery disease. PRS, polygenic risk score. LDL-C, LDL-cholesterol. TG, triglycerides.

The PRSs consisted of six million markers and explained 5.3% (adjusted *r*^*2*^) of variation in LDL-C and 4.9% in TG. In FINRISK, median LDL-C was 2.83 (95% CI 2.79-2.89) mmol/l in the lowest and 3.80 (3.72-3.88) mmol/l in the highest 5% of the LDL-C PRS distribution (Figure 1 a). Similarly, median TG was 0.99 (95% CI 0.95-1.01) mmol/l in the lowest and 1.52 (1.48-1.58) mmol/l in the highest 5% of the TG PRS distribution (Figure 1 b). The correlation between the LDL-C PRS and the TG PRS was low (*r* = 0.13). All in all, the LDL-C and TG PRSs were specific to and had clear impact on their respective lipid levels.

**Figure 1.**
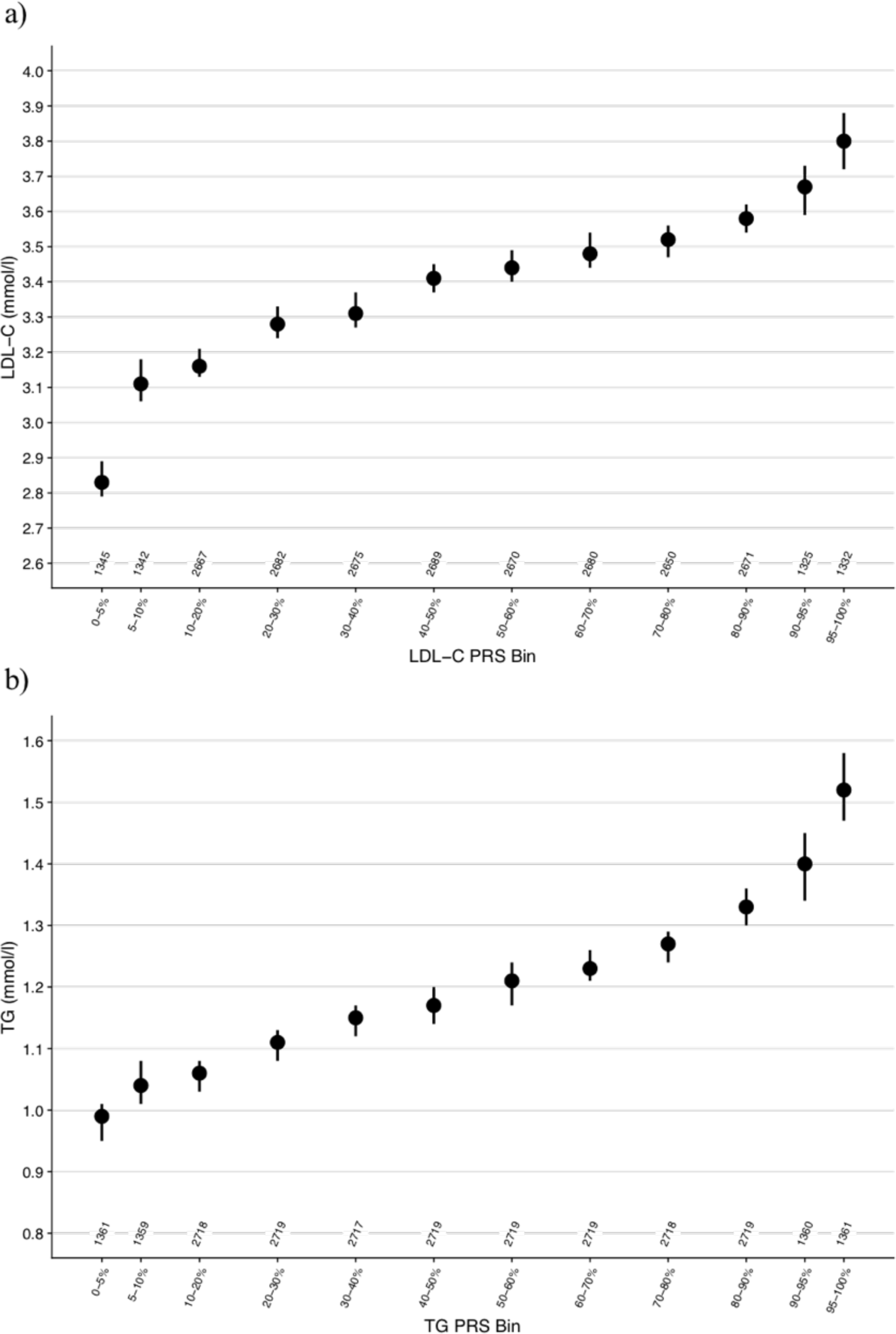
Median LDL-C (a) and TG (b) levels across the distributions of the respective PRSs in the FINRISK cohort. Numbers of individuals in the PRS bins are reported. Vertical lines represent 95% CIs. PRS, polygenic risk score. LDL-C, LDL-cholesterol. TG, triglycerides. CI, confidence interval.

### Polygenic Hyperlipidemias and CAD Risk

To assess how polygenic hyperlipidemia associates with CAD risk, we analysed 135 300 individuals including 13 695 registry-based CAD cases from the Finnish FinnGen project (Table 1). For polygenic hypercholesterolemia, CAD risk was 1.3-fold (OR 1.30 [95% CI 1.21-1.39]) higher for those in the highest 10% and 1.4-fold (OR 1.36 [95% CI 1.24-1.49]) higher for those in the highest 5% of the LDL-C PRS, compared to the remainder of the population (Figure 2 a). CAD prevalence was accordingly 54% higher (12.7% vs 8.2%) between the highest and lowest 5% of the LDL-C PRS distribution (Figure 3 a).

**Figure 2.**
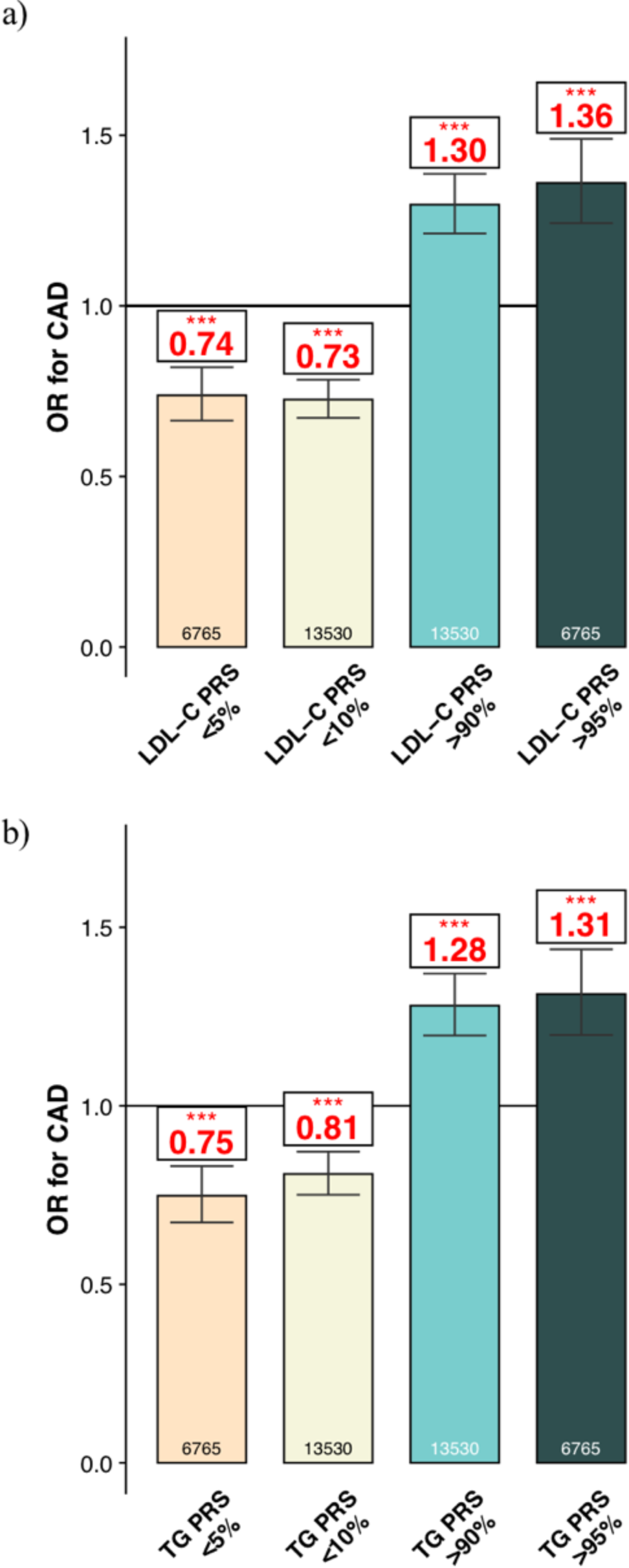
ORs for CAD across the LDL-C (a) and TG (b) PRS distributions in FinnGen. Total numbers of individuals in PRS bins are reported. ORs were estimated using logistic regression. PRS bins were compared with the remainder of the population. Error bars represent 95% CIs. PRS, polygenic risk score. LDL-C, LDL-cholesterol. CAD, coronary artery disease. OR, odds ratio. TG, triglycerides. CI, confidence interval. ‘*p* < 0.1. \**p* < 0.05. \*\**p* < 0.01. \*\*\**p* < 0.001.

**Figure 3.**
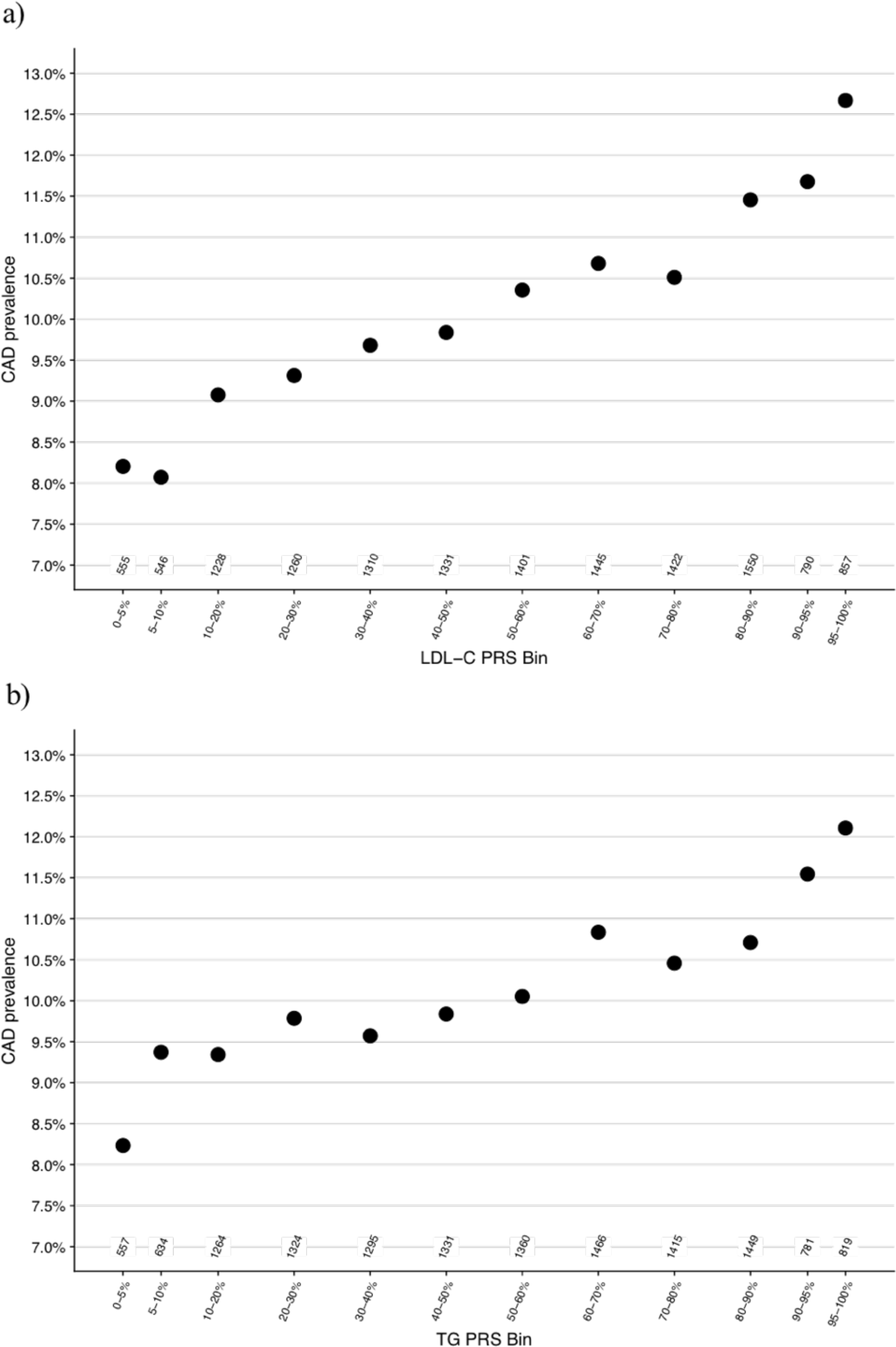
CAD prevalence across the LDL-C (a) and TG (b) PRS distributions in FinnGen. Numbers of CAD cases in PRS bins are reported. PRS, polygenic risk score. CAD, coronary artery disease. LDL-C, LDL-cholesterol. TG, triglycerides.

For polygenic hypertriglyceridemia CAD risk was 1.3-fold (OR 1.28 [95% CI 1.20-1.37]) higher for those in the highest 10% and also 1.3-fold (OR 1.31 [95% CI 1.20-1.44]) higher for those in the highest 5% of the TG PRS, compared to the remainder of the population (Figure 2 b). CAD prevalence was 47% higher (12.1% vs 8.2%) between the highest and lowest 5% of the TG PRS distribution (Figure 3 b).

We tested if the lipid PRSs improve CAD risk prediction beyond a similarly derived CAD PRS. We calculated a genome-wide CAD PRS with LDpred-based weights from a GWAS of UKBB IHD diagnoses.^21^ Comparing the highest 5% to the remainder of the population, the effects of the lipid PRSs to CAD risk were attenuated only modestly when adjusted for the CAD PRS (LDL-C PRS OR 1.26 [95% CI 1.15-1.39] and TG PRS OR 1.21 [1.10-1.32]; Figure 4).

**Figure 4.**
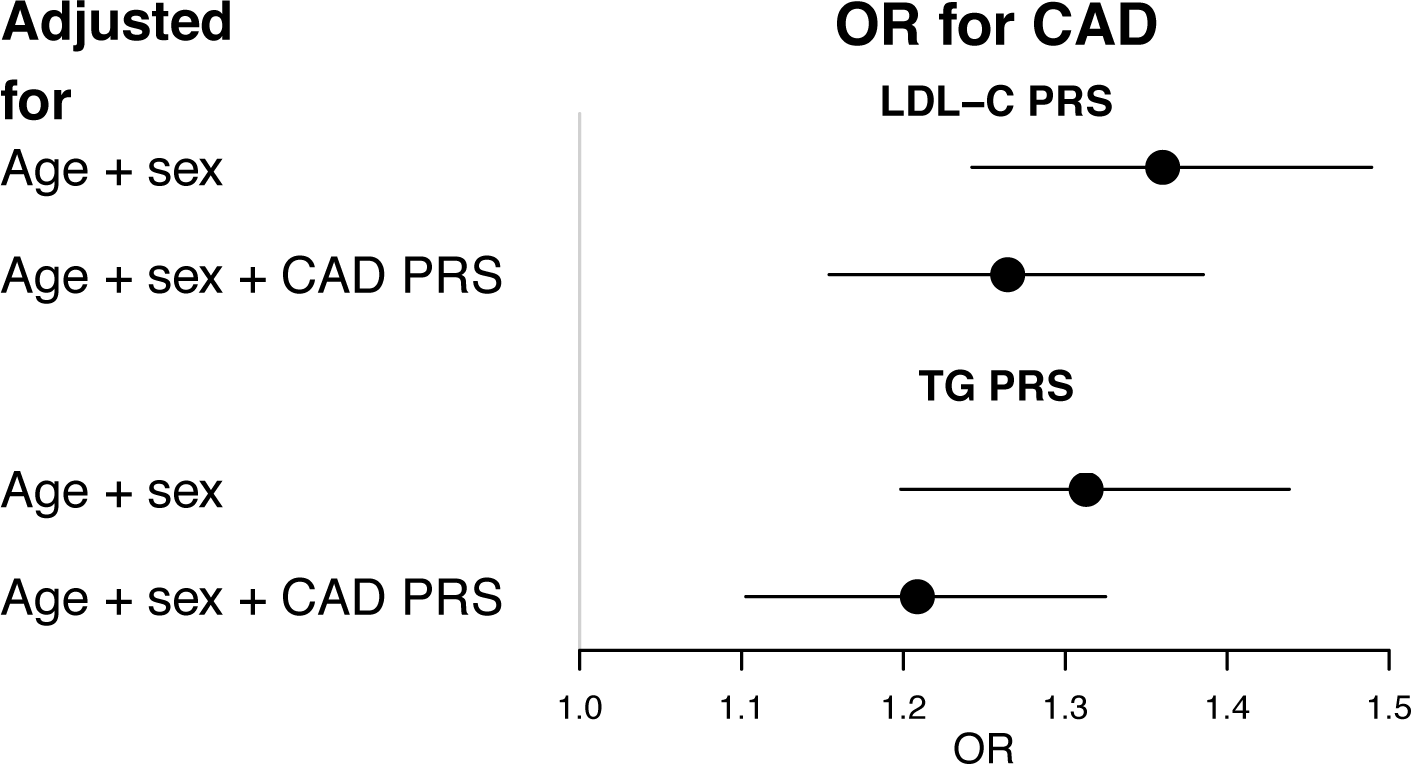
ORs for CAD for those in the highest 5% of the PRSs compared to the remainder of the population with and without adjusting for the CAD PRS in FinnGen. ORs were estimated using logistic regression. All models were additionally adjusted for age and sex. Horizontal lines represent 95% CIs. OR, odds ratio. CI, confidence interval. CAD, coronary artery disease. PRS, polygenic risk score. LDL-C, LDL-cholesterol. TG, triglycerides.

## Discussion

By developing genome-wide PRSs for LDL-C and TG, we evaluated the impact of high genetic risk for these established and causal risk factors of CAD. We showed that high polygenic burden for both LDL-C or TG associated with considerably increased LDL-C and TG levels, respectively. Similarly, polygenic hypercholesterolemia and -triglyceridemia associated with significantly increased CAD risk. Furthermore, PRSs for LDL-C and TG were mostly independent of a PRS for CAD.

Polygenic hypercholesterolemia, in our study, demonstrated 0.43 mmol/l higher LDL-C levels and 36% higher CAD risk in the highest 5% of the LDL-C PRS compared to the remainder of the population. This is considerably lower than previously reported CAD risk effects of high-impact *LDLR* FH mutations.^24^ While the established high-impact *LDLR* FH mutations directly disrupt LDL receptor function causing lifelong high LDL-C levels, the effect sizes of the individual variants contributing to polygenic hypercholesterolemia are small, and they likely increase LDL-C via multiple indirect biological pathways. Whereas monogenic FH is a severe disease with very high CAD risk, polygenic hypercholesterolemia, as captured by the current PRSs, seems to have a smaller effect on LDL-C levels and CAD risk. Furthermore, the benefit of lipid-lowering therapies in individuals with polygenic hypercholesterolemia remains unknown.

In addition to hypercholesterolemia, our results show that TG levels were 0.21 mmol/l lower in the lowest 5% of the TG PRS compared to the remainder of the population, and this translated into 25% lower CAD risk. This relationship is in line with the effect of TG-lowering loss-of-function mutations in the *APOC3* and *ANGPTL4* genes that reduce TG levels by ∼0.7-0.8 mmol/l and CAD risk by ∼40-50%..^6, 25, 26^ Our consistent results support the hypothesis that TG is causal factor for CAD. Pharmacologic TG-lowering shows promise and the benefit of TG-lowering drugs remains to be tested in individuals with polygenic hypertriglyceridemia.^11^

In our study, both LDL-C and TG PRSs associated with CAD risk also when adjusted for a CAD PRS. The key difference between intermediate biomarker PRSs (such as the lipid PRSs) and disease endpoint PRSs (such as a CAD PRS) is that for the biomarker PRSs, the mechanism of effect on clinical outcomes is more direct. The CAD PRS was based on a case-control setting of individuals with or without a CAD diagnosis with a risk of misclassifications, and correlates little with known risk factors, complicating its interpretation and clinical implications.^27^ In the presence of genetic information, biomarker PRSs could guide which intervention is taken to lower the CAD risk of an individual. How much this applies to other CAD risk factors than lipids remains unknown.

Our study has several limitations. First, as FINRISK participants fasted for a minimum of 4 hours before measuring lipid profiles, our association estimates may have been attenuated particularly between the TG PRS and TG levels. The association between the TG PRS and CAD risk, however, remains unaffected by this. Second, because the Friedewald formula is invalid for individuals with TG > 4.52 mmol/l, 456 (1.7%) FINRISK samples were excluded from LDL-C analyses.^14^ Third, some variants included in the lipid PRSs are not specific to their primary lipids and have residual effects on others. Excepting a negative association between the TG PRSs and HDL-C, however, the PRSs had only minor associations with other than their primary lipids (S2 Figure). Fourth, our weights for the lipid PRSs came from the UK population and were tested in the Finnish population; our results may have limited accuracy in other ethnicities. Replication and validation in other cohorts with lipid measurements and populations is warranted in the future.

In summary, the CAD risk associated with a high polygenic load for LDL-C or TG -increasing genetic variants was proportional to their impact on lipid levels. In contrast with a PRS for CAD, the lipid PRSs point to a known and directly modifiable risk factor enabling more straightforward clinical translation. As polygenic risk scores can also be measured at any point in life, they provide powerful tools for prioritising individuals for blood lipid panel screening and subsequent evidence-based intervention.

## Acknowledgements

We would like to thank Sari Kivikko, Huei-Yi Shen, and Ulla Tuomainen for management assistance. The FINRISK data used for the research were obtained from THL Biobank. We thank the THL DNA laboratory for its skilful work to produce the DNA samples used in the genotyping work, which was used in this study. Part of the genotyping was performed by the Institute for Molecular Medicine Finland Technology Centre, University of Helsinki. We thank all study participants for their generous participation in the FINRISK and UKBB studies. PR and JTR acknowledge support from the Doctoral Programme in Population Health, University of Helsinki, and JTR acknowledges support from the MD-PhD Programme, University of Helsinki. This research has been conducted using the UK Biobank Resource under application no 22627. The content is solely the responsibility of the authors and does not necessarily represent the official views of the National Institutes of Health.

## Appendices

**S1 Text. Supplemental Methods.**

**S1 Figure. LDL-C and TG Levels in FINRISK Surveys from 1992 to 2012.**

**S2 Figure. HDL-C, TG, and LDL-C levels in FINRISK Across the PRS Distributions.**

## Supplemental Files

### Supplemental Material

Supplemental_Material.pdf, Ripatti, Polygenic Hyperlipidemias and Coronary Artery Disease Risk.doc

