## Supplemental Material for "Polygenic Hyperlipidemias and Coronary Artery Disease Risk"

### S1 Text. Supplemental Methods.

### FinnGen

#### Steering Committee

| | |
|---|---|
| Aarno Palotie | Institute for Molecular Medicine Finland, HiLIFE, University of Helsinki, Finland |
| Mark Daly | Institute for Molecular Medicine Finland, HiLIFE, University of Helsinki, Finland |

**Pharmaceutical Companies**

| | |
|---|---|
| Howard Jacob | AbbVie, Chicago, IL, United States |
| Athena Matakidou | Astra Zeneca, Cambridge, United Kingdom |
| Heiko Runz | Biogen, Cambridge, MA, United States |
| Sally John | Biogen, Cambridge, MA, United States |
| Robert Plenge | Celgene, Summit, NJ, United States |
| Mark McCarthy | Genentech, San Francisco, CA, United States |
| Julie Hunkapiller | Genentech, San Francisco, CA, United States |
| Meg Ehm | GlaxoSmithKline, Brentford, United Kingdom |
| Dawn Waterworth | GlaxoSmithKline, Brentford, United Kingdom |
| Caroline Fox | Merck, Kenilworth, NJ, United States |
| Anders Malarstig | Pfizer, New York, NY, United States |
| Kathy Klinger | Sanofi, Paris, France |
| Kathy Call | Sanofi, Paris, France |

**University of Helsinki & Biobanks**

| | |
|---|---|
| Tomi Mäkelä | HiLIFE, University of Helsinki, Finland, Finland |
| Jaakko Kaprio | Institute for Molecular Medicine Finland, HiLIFE, Helsinki, Finland, Finland |
| Petri Virolainen | Auria Biobank / University of Turku / Hospital District of Southwest Finland, Turku, Finland |

| | |
|---|---|
| Kari Pulkki | Auria Biobank / University of Turku / Hospital District of Southwest Finland, Turku, Finland |
| Terhi Kilpi | THL Biobank / The National Institute of Health and Welfare Helsinki, Finland |
| Markus Perola | THL Biobank / The National Institute of Health and Welfare Helsinki, Finland |
| Jukka Partanen | Finnish Red Cross Blood Service / Finnish Hematology Registry and Clinical Biobank, Helsinki, Finland |
| Anne Pitkäranta | Hospital District of Helsinki and Uusimaa, Helsinki, Finland |
| Riitta Kaarteenaho | Northern Finland Biobank Borealis / University of Oulu / Northern Ostrobothnia Hospital District, Oulu, Finland |
| Seppo Vainio | Northern Finland Biobank Borealis / University of Oulu / Northern Ostrobothnia Hospital District, Oulu, Finland |
| Kimmo Savinainen | Finnish Clinical Biobank Tampere / University of Tampere / Pirkanmaa Hospital District, Tampere, Finland |
| Veli-Matti Kosma | Biobank of Eastern Finland / University of Eastern Finland / Northern Savo Hospital District, Kuopio, Finland |
| Urho Kujala | Central Finland Biobank / University of Jyväskylä / Central Finland Health Care District, Jyväskylä, Finland |

**Other Experts/Non-Voting Members**

| | |
|---|---|
| Outi Tuovila | Business Finland, Helsinki, Finland |
| Minna Hendolin | Business Finland, Helsinki, Finland |
| Raimo Pakkanen | Business Finland, Helsinki, Finland |

### Scientific Committee

**Pharmaceutical Companies**

| | |
|---|---|
| Jeff Waring | AbbVie, Chicago, IL, United States |
| Bridget Riley-Gillis | AbbVie, Chicago, IL, United States |
| Athena Matakidou | Astra Zeneca, Cambridge, United Kingdom |

| | |
|---|---|
| Heiko Runz | Biogen, Cambridge, MA, United States |
| Jimmy Liu | Biogen, Cambridge, MA, United States |
| Shameek Biswas | Celgene, Summit, NJ, United States |
| Julie Hunkapiller | Genentech, San Francisco, CA, United States |
| Dawn Waterworth | GlaxoSmithKline, Brentford, United Kingdom |
| Meg Ehm | GlaxoSmithKline, Brentford, United Kingdom |
| Dorothee Diogo | Merck, Kenilworth, NJ, United States |
| Caroline Fox | Merck, Kenilworth, NJ, United States |
| Anders Malarstig | Pfizer, New York, NY, United States |
| Catherine Marshall | Pfizer, New York, NY, United States |
| Xinli Hu | Pfizer, New York, NY, United States |
| Kathy Call | Sanofi, Paris, France |
| Kathy Klinger | Sanofi, Paris, France |
| Matthias Gossel | Sanofi, Paris, France |

**University of Helsinki & Biobanks**

| | |
|---|---|
| Samuli Ripatti | Institute for Molecular Medicine Finland, HiLIFE, Helsinki, Finland |
| Johanna Schleutker | Auria Biobank / Univ. of Turku / Hospital District of Southwest Finland, Turku, Finland |
| Markus Perola | THL Biobank / The National Institute of Health and Welfare Helsinki, Finland |
| Mikko Arvas | Finnish Red Cross Blood Service / Finnish Hematology Registry and Clinical Biobank, Helsinki, Finland |
| Olli Carpen | Hospital District of Helsinki and Uusimaa, Helsinki, Finland |
| Reetta Hinttala | Northern Finland Biobank Borealis / University of Oulu / Northern Ostrobothnia Hospital District, Oulu, Finland |
| Johannes Kettunen | Northern Finland Biobank Borealis / University of Oulu / Northern Ostrobothnia Hospital District, Oulu, Finland |

| | |
|---|---|
| Reijo Laaksonen | Finnish Clinical Biobank Tampere / University of Tampere / Pirkanmaa Hospital District, Tampere, Finland |
| Arto Mannermaa | Biobank of Eastern Finland / University of Eastern Finland / Northern Savo Hospital District, Kuopio, Finland |
| Juha Paloneva | Central Finland Biobank / University of Jyväskylä / Central Finland Health Care District, Jyväskylä, Finland |
| Urho Kujala | Central Finland Biobank / University of Jyväskylä / Central Finland Health Care District, Jyväskylä, Finland |

**Other Experts/Non-Voting Members**

| | |
|---|---|
| Outi Tuovila | Business Finland, Helsinki, Finland |
| Minna Hendolin | Business Finland, Helsinki, Finland |
| Raimo Pakkanen | Business Finland, Helsinki, Finland |

### Clinical Groups

**Neurology Group**

| | |
|---|---|
| Hilkka Soininen | Northern Savo Hospital District, Kuopio, Finland |
| Valtteri Julkunen | Northern Savo Hospital District, Kuopio, Finland |
| Anne Remes | Northern Ostrobothnia Hospital District, Oulu, Finland |
| Reetta Kälviäinen | Northern Savo Hospital District, Kuopio, Finland |
| Mikko Hiltunen | Northern Savo Hospital District, Kuopio, Finland |
| Jukka Peltola | Pirkanmaa Hospital District, Tampere, Finland |
| Pentti Tienari | Hospital District of Helsinki and Uusimaa, Helsinki, Finland |
| Juha Rinne | Hospital District of Southwest Finland, Turku, Finland |
| Adam Ziemann | AbbVie, Chicago, IL, United States |
| Jeffrey Waring | AbbVie, Chicago, IL, United States |
| Sahar Esmaeeli | AbbVie, Chicago, IL, United States |
| Nizar Smaoui | AbbVie, Chicago, IL, United States |

| | |
|---|---|
| Anne Lehtonen | AbbVie, Chicago, IL, United States |
| Susan Eaton | Biogen, Cambridge, MA, United States |
| Heiko Runz | Biogen, Cambridge, MA, United States |
| Sanni Lahdenperä | Biogen, Cambridge, MA, United States |
| Shameek Biswas | Celgene, Summit, NJ, United States |
| John Michon | Genentech, San Francisco, CA, United States |
| Geoff Kerchner | Genentech, San Francisco, CA, United States |
| Julie Hunkapiller | Genentech, San Francisco, CA, United States |
| Natalie Bowers | Genentech, San Francisco, CA, United States |
| Edmond Teng | Genentech, San Francisco, CA, United States |
| John Eicher | Merck, Kenilworth, NJ, United States |
| Vinay Mehta | Merck, Kenilworth, NJ, United States |
| Padhraig Gormley | Merck, Kenilworth, NJ, United States |
| Kari Linden | Pfizer, New York, NY, United States |
| Christopher Whelan | Pfizer, New York, NY, United States |
| Fanli Xu | GlaxoSmithKline, Brentford, United Kingdom |
| David Pulford | GlaxoSmithKline, Brentford, United Kingdom |

**Gastroenterology Group**

| | |
|---|---|
| Martti Färkkilä | Hospital District of Helsinki and Uusimaa, Helsinki, Finland |
| Sampsa Pikkarainen | Hospital District of Helsinki and Uusimaa, Helsinki, Finland |
| Airi Jussila | Pirkanmaa Hospital District, Tampere, Finland |
| Timo Blomster | Northern Ostrobothnia Hospital District, Oulu, Finland |
| Mikko Kiviniemi | Northern Savo Hospital District, Kuopio, Finland |
| Markku Voutilainen | Hospital District of Southwest Finland, Turku, Finland |
| Bob Georgantas | AbbVie, Chicago, IL, United States |
| Graham Heap | AbbVie, Chicago, IL, United States |
| Jeffrey Waring | AbbVie, Chicago, IL, United States |

| | |
|---|---|
| Nizar Smaoui | AbbVie, Chicago, IL, United States |
| Fedik Rahimov | AbbVie, Chicago, IL, United States |
| Anne Lehtonen | AbbVie, Chicago, IL, United States |
| Keith Usiskin | Celgene, Summit, NJ, United States |
| Joseph Maranville | Celgene, Summit, NJ, United States |
| Tim Lu | Genentech, San Francisco, CA, United States |
| Natalie Bowers | Genentech, San Francisco, CA, United States |
| Danny Oh | Genentech, San Francisco, CA, United States |
| John Michon | Genentech, San Francisco, CA, United States |
| Vinay Mehta | Merck, Kenilworth, NJ, United States |
| Kirsi Kalpala | Pfizer, New York, NY, United States |
| Melissa Miller | Pfizer, New York, NY, United States |
| Xinli Hu | Pfizer, New York, NY, United States |
| Linda McCarthy | GlaxoSmithKline, Brentford, United Kingdom |


**Rheumatology Group**


| | |
|---|---|
| Kari Eklund | Hospital District of Helsinki and Uusimaa, Helsinki, Finland |
| Antti Palomäki | Hospital District of Southwest Finland, Turku, Finland |
| Pia Isomäki | Pirkanmaa Hospital District, Tampere, Finland |
| Laura Pirilä | Hospital District of Southwest Finland, Turku, Finland |
| Oili Kaipiainen-Seppänen | Northern Savo Hospital District, Kuopio, Finland |
| Johanna Huhtakangas | Northern Ostrobothnia Hospital District, Oulu, Finland |
| Bob Georgantas | AbbVie, Chicago, IL, United States |
| Jeffrey Waring | AbbVie, Chicago, IL, United States |
| Fedik Rahimov | AbbVie, Chicago, IL, United States |
| Apinya Lertratanakul | AbbVie, Chicago, IL, United States |
| Nizar Smaoui | AbbVie, Chicago, IL, United States |

| | |
|---|---|
| Anne Lehtonen | AbbVie, Chicago, IL, United States |
| David Close | Astra Zeneca, Cambridge, United Kingdom |
| Marla Hochfeld | Celgene, Summit, NJ, United States |
| Natalie Bowers | Genentech, San Francisco, CA, United States |
| John Michon | Genentech, San Francisco, CA, United States |
| Dorothee Diogo | Merck, Kenilworth, NJ, United States |
| Vinay Mehta | Merck, Kenilworth, NJ, United States |
| Kirsi Kalpala | Pfizer, New York, NY, United States |
| Nan Bing | Pfizer, New York, NY, United States |
| Xinli Hu | Pfizer, New York, NY, United States |
| Jorge Esparza Gordillo | GlaxoSmithKline, Brentford, United Kingdom |
| Nina Mars | Institute for Molecular Medicine Finland, HiLIFE, Helsinki, Finland |

**Pulmonology Group**

| | |
|---|---|
| Tarja Laitinen | Pirkanmaa Hospital District, Tampere, Finland |
| Margit Pelkonen | Northern Savo Hospital District, Kuopio, Finland |
| Paula Kauppi | Hospital District of Helsinki and Uusimaa, Helsinki, Finland |
| Hannu Kankaanranta | Pirkanmaa Hospital District, Tampere, Finland |
| Terttu Harju | Northern Ostrobothnia Hospital District, Oulu, Finland |
| Nizar Smaoui | AbbVie, Chicago, IL, United States |
| David Close | Astra Zeneca, Cambridge, United Kingdom |
| Steven Greenberg | Celgene, Summit, NJ, United States |
| Hubert Chen | Genentech, San Francisco, CA, United States |
| Natalie Bowers | Genentech, San Francisco, CA, United States |
| John Michon | Genentech, San Francisco, CA, United States |
| Vinay Mehta | Merck, Kenilworth, NJ, United States |
| Jo Betts | GlaxoSmithKline, Brentford, United Kingdom |
| Soumitra Ghosh | GlaxoSmithKline, Brentford, United Kingdom |

**Cardiometabolic Diseases Group**

| | |
|---|---|
| Veikko Salomaa | The National Institute of Health and Welfare Helsinki, Finland |
| Teemu Niiranen | The National Institute of Health and Welfare Helsinki, Finland |
| Markus Juonala | Hospital District of Southwest Finland, Turku, Finland |
| Kaj Metsärinne | Hospital District of Southwest Finland, Turku, Finland |
| Mika Kähönen | Pirkanmaa Hospital District, Tampere, Finland |
| Juhani Junttila | Northern Ostrobothnia Hospital District, Oulu, Finland |
| Markku Laakso | Northern Savo Hospital District, Kuopio, Finland |
| Jussi Pihlajamäki | Northern Savo Hospital District, Kuopio, Finland |
| Juha Sinisalo | Hospital District of Helsinki and Uusimaa, Helsinki, Finland |
| Marja-Riitta Taskinen | Hospital District of Helsinki and Uusimaa, Helsinki, Finland |
| Tiinamaija Tuomi | Hospital District of Helsinki and Uusimaa, Helsinki, Finland |
| Jari Laukkanen | Central Finland Health Care District, Jyväskylä, Finland |
| Ben Challis | Astra Zeneca, Cambridge, United Kingdom |
| Andrew Peterson | Genentech, San Francisco, CA, United States |
| Julie Hunkapiller | Genentech, San Francisco, CA, United States |
| Natalie Bowers | Genentech, San Francisco, CA, United States |
| John Michon | Genentech, San Francisco, CA, United States |
| Dorothee Diogo | Merck, Kenilworth, NJ, United States |
| Audrey Chu | Merck, Kenilworth, NJ, United States |
| Vinay Mehta | Merck, Kenilworth, NJ, United States |
| Jaakko Parkkinen | Pfizer, New York, NY, United States |
| Melissa Miller | Pfizer, New York, NY, United States |
| Anthony Muslin | Sanofi, Paris, France |
| Dawn Waterworth | GlaxoSmithKline, Brentford, United Kingdom |

**Oncology Group**

| | |
|---|---|
| Heikki Joensuu | Hospital District of Helsinki and Uusimaa, Helsinki, Finland |
| Tuomo Meretoja | Hospital District of Helsinki and Uusimaa, Helsinki, Finland |

| | |
|---|---|
| Olli Carpen | Hospital District of Helsinki and Uusimaa, Helsinki, Finland |
| Lauri Aaltonen | Hospital District of Helsinki and Uusimaa, Helsinki, Finland |
| Annika Auranen | Pirkanmaa Hospital District , Tampere, Finland |
| Peeter Karihtala | Northern Ostrobothnia Hospital District, Oulu, Finland |
| Saila Kauppila | Northern Ostrobothnia Hospital District, Oulu, Finland |
| Päivi Auvinen | Northern Savo Hospital District, Kuopio, Finland |
| Klaus Elenius | Hospital District of Southwest Finland, Turku, Finland |
| Relja Popovic | AbbVie, Chicago, IL, United States |
| Jeffrey Waring | AbbVie, Chicago, IL, United States |
| Bridget Riley-Gillis | AbbVie, Chicago, IL, United States |
| Anne Lehtonen | AbbVie, Chicago, IL, United States |
| Athena Matakidou | Astra Zeneca, Cambridge, United Kingdom |
| Jennifer Schutzman | Genentech, San Francisco, CA, United States |
| Julie Hunkapiller | Genentech, San Francisco, CA, United States |
| Natalie Bowers | Genentech, San Francisco, CA, United States |
| John Michon | Genentech, San Francisco, CA, United States |
| Vinay Mehta | Merck, Kenilworth, NJ, United States |
| Andrey Loboda | Merck, Kenilworth, NJ, United States |
| Aparna Chhibber | Merck, Kenilworth, NJ, United States |
| Heli Lehtonen | Pfizer, New York, NY, United States |
| Stefan McDonough | Pfizer, New York, NY, United States |
| Marika Crohns | Sanofi, Paris, France |
| Diptee Kulkarni | GlaxoSmithKline, Brentford, United Kingdom |

**Opthalmology Group**

| | |
|---|---|
| Kai Kaarniranta | Northern Savo Hospital District, Kuopio, Finland |
| Joni Turunen | Hospital District of Helsinki and Uusimaa, Helsinki, Finland |
| Terhi Ollila | Hospital District of Helsinki and Uusimaa, Helsinki, Finland |

| | |
|---|---|
| Sanna Seitsonen | Hospital District of Helsinki and Uusimaa, Helsinki, Finland |
| Hannu Uusitalo | Pirkanmaa Hospital District, Tampere, Finland |
| Vesa Aaltonen | Hospital District of Southwest Finland, Turku, Finland |
| Hannele Uusitalo-Järvinen | Pirkanmaa Hospital District, Tampere, Finland |
| Marja Luodonpää | Northern Ostrobothnia Hospital District, Oulu, Finland |
| Nina Hautala | Northern Ostrobothnia Hospital District, Oulu, Finland |
| Heiko Runz | Biogen, Cambridge, MA, United States |
| Erich Strauss | Genentech, San Francisco, CA, United States |
| Natalie Bowers | Genentech, San Francisco, CA, United States |
| Hao Chen | Genentech, San Francisco, CA, United States |
| John Michon | Genentech, San Francisco, CA, United States |
| Anna Podgornaia | Merck, Kenilworth, NJ, United States |
| Vinay Mehta | Merck, Kenilworth, NJ, United States |
| Dorothee Diogo | Merck, Kenilworth, NJ, United States |
| Joshua Hoffman | GlaxoSmithKline, Brentford, United Kingdom |

**Dermatology Group**

| | |
|---|---|
| Kaisa Tasanen | Northern Ostrobothnia Hospital District, Oulu, Finland |
| Laura Huilaja | Northern Ostrobothnia Hospital District, Oulu, Finland |
| Katariina Hannula-Jouppi | Hospital District of Helsinki and Uusimaa, Helsinki, Finland |
| Teea Salmi | Pirkanmaa Hospital District, Tampere, Finland |
| Sirkku Peltonen | Hospital District of Southwest Finland, Turku, Finland |
| Leena Koulu | Hospital District of Southwest Finland, Turku, Finland |
| Ilkka Harvima | Northern Savo Hospital District, Kuopio, Finland |
| Kirsi Kalpala | Pfizer, New York, NY, United States |
| Ying Wu | Pfizer, New York, NY, United States |

| | |
|---|---|
| David Choy | Genentech, San Francisco, CA, United States |
| John Michon | Genentech, San Francisco, CA, United States |
| Nizar Smaoui | AbbVie, Chicago, IL, United States |
| Fedik Rahimov | AbbVie, Chicago, IL, United States |
| Anne Lehtonen | AbbVie, Chicago, IL, United States |
| Dawn Waterworth | GlaxoSmithKline, Brentford, United Kingdom |

### FinnGen Teams

#### Administration Team

| | |
|---|---|
| Anu Jalanko | Institute for Molecular Medicine Finland, HiLIFE, University of Helsinki, Finland |
| Risto Kajanne | Institute for Molecular Medicine Finland, HiLIFE, University of Helsinki, Finland |
| Ulrike Lyhs | Institute for Molecular Medicine Finland, HiLIFE, University of Helsinki, Finland |

#### Communication

| | |
|---|---|
| Mari Kaunisto | Institute for Molecular Medicine Finland, HiLIFE, University of Helsinki, Finland |

#### Analysis Team

| | |
|---|---|
| Justin Wade Davis | AbbVie, Chicago, IL, United States |
| Bridget Riley-Gillis | AbbVie, Chicago, IL, United States |
| Danjuma Quarless | AbbVie, Chicago, IL, United States |
| Slavé Petrovski | Astra Zeneca, Cambridge, United Kingdom |
| Jimmy Liu | Biogen, Cambridge, MA, United States |
| Chia-Yen Chen | Biogen, Cambridge, MA, United States |
| Paola Bronson | Biogen, Cambridge, MA, United States |
| Robert Yang | Celgene, Summit, NJ, United States |
| Joseph Maranville | Celgene, Summit, NJ, United States |
| Shameek Biswas | Celgene, Summit, NJ, United States |
| Diana Chang | Genentech, San Francisco, CA, United States |
| Julie Hunkapiller | Genentech, San Francisco, CA, United States |

| | |
|---|---|
| Tushar Bhangale | Genentech, San Francisco, CA, United States |
| Natalie Bowers | Genentech, San Francisco, CA, United States |
| Dorothee Diogo | Merck, Kenilworth, NJ, United States |
| Emily Holzinger | Merck, Kenilworth, NJ, United States |
| Padhraig Gormley | Merck, Kenilworth, NJ, United States |
| Xulong Wang | Merck, Kenilworth, NJ, United States |
| Xing Chen | Pfizer, New York, NY, United States |
| Åsa Hedman | Pfizer, New York, NY, United States |
| Kirsi Auro | GlaxoSmithKline, Brentford, United Kingdom |
| Clarence Wang | Sanofi, Paris, France |
| Ethan Xu | Sanofi, Paris, France |
| Franck Auge | Sanofi, Paris, France |
| Clement Chatelain | Sanofi, Paris, France |
| Mitja Kurki | Institute for Molecular Medicine Finland, HiLIFE, University of Helsinki, Finland / Broad Institute, Cambridge, MA, United States |
| Samuli Ripatti | Institute for Molecular Medicine Finland, HiLIFE, University of Helsinki, Finland |
| Mark Daly | Institute for Molecular Medicine Finland, HiLIFE, University of Helsinki, Finland |
| Juha Karjalainen | Institute for Molecular Medicine Finland, HiLIFE, University of Helsinki, Finland / Broad Institute, Cambridge, MA, United States |
| Aki Havulinna | Institute for Molecular Medicine Finland, HiLIFE, University of Helsinki, Finland |
| Anu Jalanko | Institute for Molecular Medicine Finland, HiLIFE, University of Helsinki, Finland |
| Kimmo Palin | University of Helsinki, Helsinki, Finland |
| Priit Palta | Institute for Molecular Medicine Finland, HiLIFE, University of Helsinki, Finland |
| Pietro Della Briotta | |
| Parolo | Institute for Molecular Medicine Finland, HiLIFE, University of Helsinki, Finland |
| Wei Zhou | Broad Institute, Cambridge, MA, United States |
| Susanna Lemmelä | Institute for Molecular Medicine Finland, HiLIFE, University of Helsinki, Finland |
| Manuel Rivas | University of Stanford, Stanford, CA, United States |

| | |
|---|---|
| Jarmo Harju | Institute for Molecular Medicine Finland, HiLIFE, University of Helsinki, Finland |
| Aarno Palotie | Institute for Molecular Medicine Finland, HiLIFE, University of Helsinki, Finland |
| Arto Lehisto | Institute for Molecular Medicine Finland, HiLIFE, University of Helsinki, Finland |
| Andrea Ganna | Institute for Molecular Medicine Finland, HiLIFE, University of Helsinki, Finland |
| Vincent Llorens | Institute for Molecular Medicine Finland, HiLIFE, University of Helsinki, Finland |
| Antti Karlsson | Auria Biobank / Univ. of Turku / Hospital District of Southwest Finland, Turku, Finland |
| Kati Kristiansson | THL Biobank / The National Institute of Health and Welfare Helsinki, Finland |
| Mikko Arvas | Finnish Red Cross Blood Service / Finnish Hematology Registry and Clinical Biobank, Helsinki, Finland |
| Kati Hyvärinen | Finnish Red Cross Blood Service / Finnish Hematology Registry and Clinical Biobank, Helsinki, Finland |
| Jarmo Ritari | Finnish Red Cross Blood Service / Finnish Hematology Registry and Clinical Biobank, Helsinki, Finland |
| Tiina Wahlfors | Finnish Red Cross Blood Service / Finnish Hematology Registry and Clinical Biobank, Helsinki, Finland |
| Miika Koskinen | Hospital District of Helsinki and Uusimaa, Helsinki, Finland BB/HUS/Univ Hosp Districts |
| Olli Carpen | Hospital District of Helsinki and Uusimaa, Helsinki, Finland BB/HUS/Univ Hosp Districts |
| Johannes Kettunen | Northern Finland Biobank Borealis / University of Oulu / Northern Ostrobothnia Hospital District, Oulu, Finland |
| Katri Pylkäs | Northern Finland Biobank Borealis / University of Oulu / Northern Ostrobothnia Hospital District, Oulu, Finland |
| Marita Kalaoja | Northern Finland Biobank Borealis / University of Oulu / Northern Ostrobothnia Hospital District, Oulu, Finland |
| Minna Karjalainen | Northern Finland Biobank Borealis / University of Oulu / Northern Ostrobothnia Hospital District, Oulu, Finland |

| | |
|---|---|
| Tuomo Mantere | Northern Finland Biobank Borealis / University of Oulu / Northern Ostrobothnia Hospital District, Oulu, Finland |
| Eeva Kangasniemi | Finnish Clinical Biobank Tampere / University of Tampere / Pirkanmaa Hospital District, Tampere, Finland |
| Sami Heikkinen | Biobank of Eastern Finland / University of Eastern Finland / Northern Savo Hospital District, Kuopio, Finland |
| Arto Mannermaa | Biobank of Eastern Finland / University of Eastern Finland / Northern Savo Hospital District, Kuopio, Finland |
| Eija Laakkonen | Central Finland Biobank / University of Jyväskylä / Central Finland Health Care District, Jyväskylä, Finland |
| Juha Kononen | Central Finland Biobank / University of Jyväskylä / Central Finland Health Care District, Jyväskylä, Finland |

**Sample Collection Coordination**

| | |
|---|---|
| Anu Loukola | Hospital District of Helsinki and Uusimaa, Helsinki, Finland |

**Sample Logistics**

| | |
|---|---|
| Päivi Laiho | THL Biobank / The National Institute of Health and Welfare Helsinki, Finland |
| Tuuli Sistonen | THL Biobank / The National Institute of Health and Welfare Helsinki, Finland |
| Essi Kaiharju | THL Biobank / The National Institute of Health and Welfare Helsinki, Finland |
| Markku Laukkanen | THL Biobank / The National Institute of Health and Welfare Helsinki, Finland |
| Elina Järvensivu | THL Biobank / The National Institute of Health and Welfare Helsinki, Finland |
| Sini Lähteenmäki | THL Biobank / The National Institute of Health and Welfare Helsinki, Finland |
| Lotta Männikkö | THL Biobank / The National Institute of Health and Welfare Helsinki, Finland |
| Regis Wong | THL Biobank / The National Institute of Health and Welfare Helsinki, Finland |

**Registry Data Operations**

| | |
|---|---|
| Kati Kristiansson | THL Biobank / The National Institute of Health and Welfare Helsinki, Finland |
| Hannele Mattsson | THL Biobank / The National Institute of Health and Welfare Helsinki, Finland |

| | |
|---|---|
| Susanna Lemmelä | Institute for Molecular Medicine Finland, HiLIFE, University of Helsinki, Finland |
| Tero Hiekkalinna | THL Biobank / The National Institute of Health and Welfare Helsinki, Finland |
| Manuel González Jiménez | THL Biobank / The National Institute of Health and Welfare Helsinki, Finland |

**Genotyping**

| | |
|---|---|
| Kati Donner | Institute for Molecular Medicine Finland, HiLIFE, University of Helsinki, Finland |

**Sequencing Informatics**

| | |
|---|---|
| Priit Palta | Institute for Molecular Medicine Finland, HiLIFE, University of Helsinki, Finland |
| Kalle Pärn | Institute for Molecular Medicine Finland, HiLIFE, University of Helsinki, Finland |
| Javier Nunez-Fontarnau | Institute for Molecular Medicine Finland, HiLIFE, University of Helsinki, Finland |

**Data Management and IT Infrastructure**

| | |
|---|---|
| Jarmo Harju | Institute for Molecular Medicine Finland, HiLIFE, University of Helsinki, Finland |
| Elina Kilpeläinen | Institute for Molecular Medicine Finland, HiLIFE, University of Helsinki, Finland |
| Timo P. Sipilä | Institute for Molecular Medicine Finland, HiLIFE, University of Helsinki, Finland |
| Georg Brein | Institute for Molecular Medicine Finland, HiLIFE, University of Helsinki, Finland |
| Alexander Dada | Institute for Molecular Medicine Finland, HiLIFE, University of Helsinki, Finland |
| Ghazal Awaisa | Institute for Molecular Medicine Finland, HiLIFE, University of Helsinki, Finland |
| Anastasia Shcherban | Institute for Molecular Medicine Finland, HiLIFE, University of Helsinki, Finland |
| Tuomas Sipilä | Institute for Molecular Medicine Finland, HiLIFE, University of Helsinki, Finland |

**Clinical Endpoint Development**

| | |
|---|---|
| Hannele Laivuori | Institute for Molecular Medicine Finland, HiLIFE, University of Helsinki, Finland |
| Aki Havulinna | Institute for Molecular Medicine Finland, HiLIFE, University of Helsinki, Finland |
| Susanna Lemmelä | Institute for Molecular Medicine Finland, HiLIFE, University of Helsinki, Finland |
| Tuomo Kiiskinen | Institute for Molecular Medicine Finland, HiLIFE, University of Helsinki, Finland |

**Trajectory Team**

| | |
|---|---|
| Tarja Laitinen | Pirkanmaa Hospital District, Tampere, Finland |

| | |
|---|---|
| Harri Siirtola | University of Tampere, Tampere, Finland |
| Javier Gracia Tabuenca | University of Tampere, Tampere, Finland |

**Biobank Directors**

| | |
|---|---|
| Lila Kallio | Auria Biobank, Turku, Finland |
| Sirpa Soini | THL Biobank, Helsinki, Finland |
| Jukka Partanen | Blood Service Biobank, Helsinki, Finland |
| Kimmo Pitkänen | Helsinki Biobank, Helsinki, Finland |
| Seppo Vainio | Northern Finland Biobank Borealis, Oulu, Finland |
| Kimmo Savinainen | Tampere Biobank, Tampere, Finland |
| Veli-Matti Kosma | Biobank of Eastern Finland, Kuopio, Finland |
| Teijo Kuopio | Central Finland Biobank, Jyväskylä, Finland |

### Genotyping and Imputation

A total of 27 039 FINRISK samples were genotyped using several arrays: the HumanCoreExome BeadChip, the Human610-Quad BeadChip, the Affymetrix6.0, and the Infinium HumanOmniExpress (Illumina Inc., San Diego, CA, USA). Genotypes were called together with other available data sets using zCall at the Institute for Molecular Medicine Finland (FIMM).[1] After sample-wise quality control (QC) (exclude samples with sex discrepancies, high genotype missingness [> 5%], excess heterozygosity [±4 SD], or non-European ancestry) and variant-wise QC (exclude SNPs with high missingness [> 2%], deviation from Hardy-Weinberg equilibrium [HWE, $p < 1\times10^{-6}$], or low minor allele count [MAC, < 3]) steps, the samples were pre-phased using Eagle2 (version 2.3).[2] Genotypes were imputed using a Finnish population-specific reference panel comprising 2690 high-coverage WGS and 5092 WES samples with IMPUTE2 (version 2.3.2) that allows the usage of two panels at the same time (the "merge_ref_panels" option).[3, 4] Post-imputation QC involved excluding variants imputed with imputation INFO < 0.7.

A total of 96 499 FinnGen samples were genotyped with Illumina (Illumina Inc., San Diego, CA, USA) and Affymetrix arrays (Thermo Fisher Scientific, Santa Clara, CA, USA). Illumina genotypes were called with GenCall and zCall, and Affymetrix with genotypes with AxiomGT1.[1] After sample-wise QC (exclude samples with sex discrepancies, high genotype missingness [> 5%], excess heterozygosity [±4 SD], or non-Finnish ancestry) and variant-wise QC (exclude SNPs with high missingness [> 2%], deviation from HWE [$p < 1\times10^{-6}$], or low MAC [< 3]) steps, the samples were pre-phased using Eagle2 (version 2.3.5) with number of conditioning haplotypes set to 20 000.[2] Genotypes were imputed using the Finnish population-specific reference panel SISu v3 with Beagle (version 4.1 08Jun17.d8b) as described.[5, 6] The SISu v3 imputation reference panel was developed using high-coverage (25-30x) WGS data generated at the Broad Institute of MIT and Harvard (Broad Institute) and at the McDonnell Genome Institute at Washington University and jointly processed at the Broad Institute. The variant call set was produced using GATK HaplotypeCaller following best practices.[7] Genotype, sample, and variant-wise QC was applied in an iterative manner using Hail (version 0.1).[8] The resulting high-quality WGS data for 3775 individuals were phased with Eagle 2.3.5 as described above.[2] Post-imputation QC involved excluding variants imputed with imputation INFO < 0.7.

A total of 343 672 UKBB samples were genotyped centrally using either the custom UK BiLEVE Axiom Array (~10%) or the closely related UK Biobank Axiom Array (~90%) with > 800 000 genetic markers across the genome.[9] Genotypes were called centrally using a custom pipeline by Affymetrix. After sample-wise QC (exclude samples with sex discrepancies, high missingness, excess heterozygosity, putative sex chromosome aneuploidy, or withdrawal of consent) and variant-wise QC (batch effects, plate effects, sex effects, array effects, exclude SNPs with deviation from HWE or discordance across control replicates), the samples were pre-phased centrally using SHAPEIT3 (version 2.3.5).[9, 10] Genotypes were imputed to ~92 million markers centrally using the Haplotype Reference Consortium resource, the UK10K panel, and the 1000 Genomes panel with IPMUTE4.[11] We limited our analysis to non-related to white British ancestry individuals, with exclusion lists constructed centrally using a combination of self-reported ancestry and genetically confirmed ancestry using principal components.

**S1 Figure.**

a)

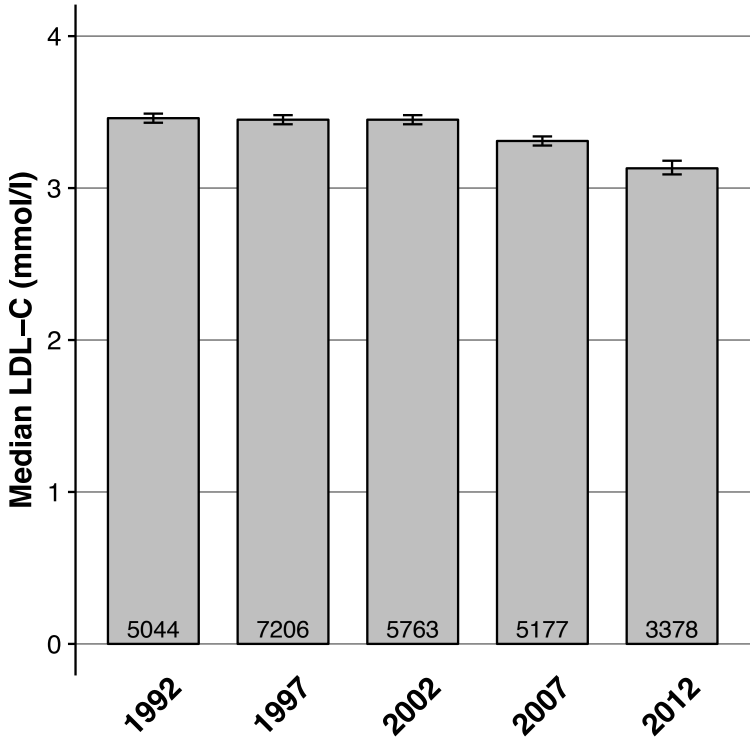

Median LDL–C (mmol/l)
0
1
2
3
4
5044
7206
5763
5177
3378
1992
1997
2002
2007
2012


b)

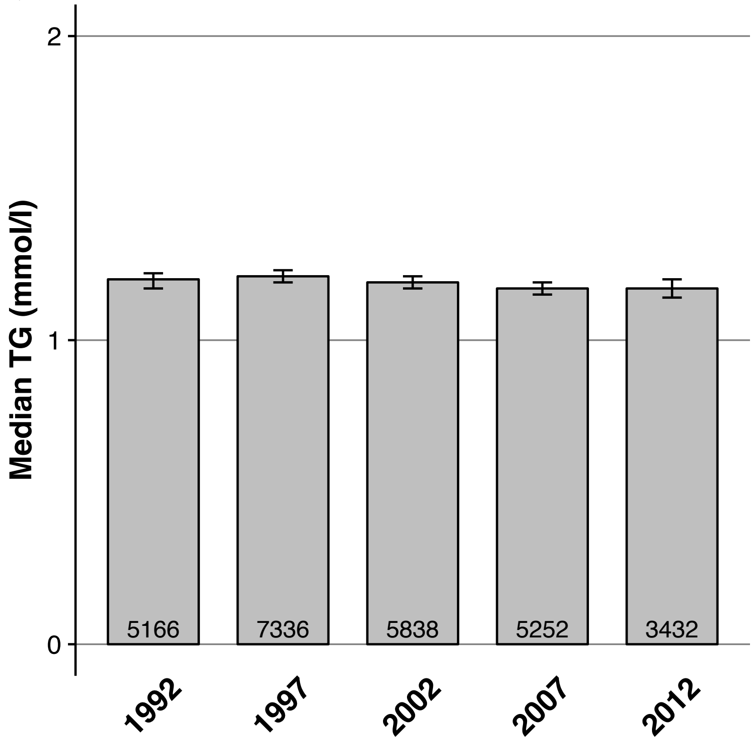

Median TG (mmol/l)
0
1
2
5166
7336
5838
5252
3432
1992
1997
2002
2007
2012

Mean LDL-C (a) and median TG (b) levels in the National FINRISK Study collection surveys from 1992, 1997, 2002, 2007, and 2012 are shown. The collections are independent, random and representative population samples of Finland. LDL-C was calculated using the Friedewald formula; the effect of lipid-lowering therapy in those using medication at the time of lipid measurement was adjusted for by dividing LDL-C by 0.7 as utilized previously.[12] Numbers of individuals in different collections are reported at the bottom of the bars. Error bars represent 95% CIs. LDL-C, LDL-cholesterol. TG, triglycerides.

**S2 Figure.**

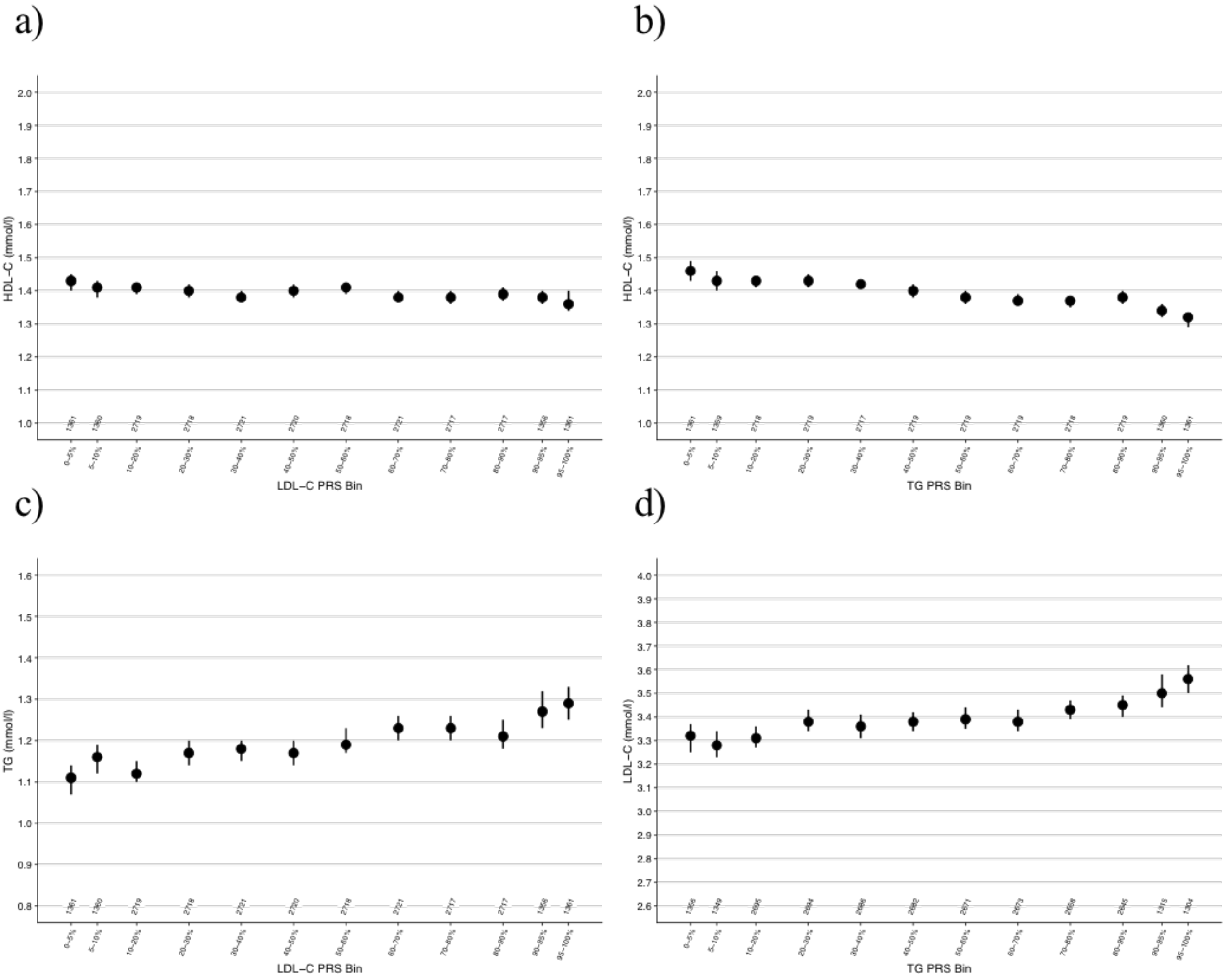
a)
HDL−C (mmol/l)
LDL−C PRS Bin
b)
HDL−C (mmol/l)
TG PRS Bin
c)
TG (mmol/l)
LDL−C PRS Bin
d)
LDL−C (mmol/l)
TG PRS Bin

Median HDL-C (a and b), TG (c), and LDL-C (d) levels across the LDL-C (a and c) and TG (b and d) PRS distributions in the FINRISK cohort. Numbers of individuals in the PRS bins are reported. Vertical lines represent 95% CIs. HDL-C, HDL-cholesterol. TG, triglycerides. LDL-C, LDL-cholesterol. PRS, polygenic risk score. CI, confidence interval.
